# Salient Safety Conditioning Improves Novel Discrimination Learning

**DOI:** 10.1101/2020.06.22.160465

**Authors:** I. Nahmoud, J. Ganay Vasquez, H. Cho, T. Dennis-Tiwary, E. Likhtik

**Affiliations:** Chemistry Dept., Hunter College, CUNY, New York, NY; Department of Psychology, The Graduate Center, CUNY, NY; Psychology Dept., Hunter College, CUNY, New York, NY; Biology Dept., Hunter College, CUNY, New York, NY; Program in Biology, The Graduate Center, CUNY, NY

**Keywords:** safety learning, fear discrimination, innate anxiety, emotion regulation, salience

## Abstract

Generalized fear is one purported mechanism of anxiety that is a target of clinical and basic research. Impaired fear discrimination has been primarily examined from the perspective of increased fear learning, rather than how learning about non-threatening stimuli affects fear discrimination. To address this question, we tested how three Safety Conditioning protocols with varied levels of salience allocated to the safety cue compared to classic Fear Conditioning in their impact on subsequent innate anxiety, and differential fear learning of new aversive and neutral cues. Using a high anxiety strain of mice (129SvEv, Taconic), we show that Fear Conditioned animals show little exploration of the anxiogenic center of an open field 24 hours later, and poor discrimination during new differential conditioning 7 days later. Three groups of mice underwent Safety Conditioning, (i) the safety tone was unpaired with a shock, (ii) the safety tone was unpaired with the shock and co-terminated with a house light signaling the end of the safety period, and (iii) the safety tone was unpaired with the shock and its beginning co-occurred with a house light, signaling the start of a safety period. Mice from all Safety Conditioning groups showed higher levels of open field exploration than the Fear Conditioned mice 24 hours after training. Furthermore, Safety Conditioned animals showed improved discrimination learning of a novel non-threat, with the Salient Beginning safety conditioned group performing best. These findings indicate that high anxiety animals benefit from salient safety training to improve exploration and discrimination of new non-threating stimuli.

**Highlights:**

- Safety training using salient cues improves safety learning in high anxiety mice
- Salient Safety training improves novel fear discrimination learning
- Safety training improves exploration in a novel anxiogenic environment

## 1. Introduction

Stress- and anxiety-related disorders are characterized by specific disruptions in fear learning and the regulation of fear, making these processes a target of cognitive behavioral therapies (1). For example, a variety of cognitive and behavioral adaptations, such as fearful or avoidant responses to stimuli that were never associated with danger, is posited to drive the emergence of overgeneralized fear responses (2–4). In keeping with this, people diagnosed with anxiety disorders show elevated fear to cues associated with aversive outcomes along with other stimuli that resemble the aversive cues (3–11). Similarly, in lab settings, fear generalization is characterized by defensive responding to stimuli that were explicitly paired with an aversive outcome (CS+) as well as to those that were never explicitly paired with anything aversive (CS−, (2–5, 7, 8, 12).However, exposure to the aversive stimulus itself poses a potent barrier to discrimination because fear conditioning widens stimulus generalization curves around the aversive stimulus, and increase the perceptual threshold needed to recognize a non-aversive stimulus as different (9–11). An intervention implication of this literature is that improving discrimination of the CS−, and identifying conditions and processes that promote its effective acquisition and retrieval, has the potential to inhibit anxiety-related fear generalization.

The CS− is a feature-negative cue, and is therefore more difficult to learn than the feature positive, aversive CS+ that drives stronger and faster acquisition (13, 14). One approach to increase CS− discrimination in high anxiety individuals, is to train anxious subjects to attend to feature-negative stimuli, thereby hoping to counteract the generalizing effects of the CS+. Safety cues are a stronger variant of the feature-negative stimulus because they inhibit fearful responding, and have rewarding and motivational properties, such as bar pressing for a reward, and stronger conditioned place preference for the area where the safety cue was presented (15–17). Safety stimulus features overlap with those of the discriminated CS−, but are a stronger variant because they signal the removal of danger whereas the CS− is simply not associated with danger, thereby making the safety cue an excellent candidate for training the behavior and networks of CS− discrimination. In the present study, first we test whether manipulating the salience of safety cues, thereby increasing attention to these feature-negative stimuli, serves to enhance their impact on fear inhibition in the conditioning context. Second, we examine whether conditioned safety and fear cues impact exploration in a novel anxiogenic environment. Third, we reasoned that salient safety cues could train the fear suppression network, and thus also improve fear suppression to the CS− during subsequent discrimination learning, resulting in less overgeneralized fear.

Safety learning and CS− discrimination are likely to have some overlap in their underlying neural processes. For example, discrimination of a safe stimulus is modulated in the bed nucleus of the stria terminalis (BNST), likely via regulation of the 5HT-2C receptors (18–21). In addition, safety learning drives firing in the basolateral amygdala (BLA), with a subset of safety-cue activated cells also driven by cues signaling reward (22). The BLA also differentially encodes fear-associated and non-associated stimuli with different subsets of cells active during retrieval of the CS+ and the CS− (22–26). Safety learning also upregulates activity in the medial prefrontal cortex (mPFC), an area that is associated with suppression of fearful responses during discrimination of non-threat and fear extinction learning (21, 27–29). In keeping with this, mPFC activity promotes encoding of the CS+ and the CS− during discrimination, and damage to the mPFC in animals impairs fear discrimination learning (30, 31). Similarly, in human anxiety, disrupted activity in the mPFC is associated with fear generalization (32, 33). Moreover, BLA communication with the mPFC is key for discrimination learning and retrieval in rodents and non-human primates, when BLA to mPFC dominates early in acquisition, whereas mPFC to BLA signaling is prevalent during CS− discrimination (8, 34–36). Imaging work in humans suggest that a similar increase in mPFC activity and a dampening of the amygdala is observed during safety cue retrieval as well (37).

Drawing on prior research, we reasoned that safety learning may engage some of the same regions and communication patterns that encode the CS−, thereby improving subsequent discrimination learning, and decreasing generalized anxiety. To examine this, we used the129SvEv strain of mice because we and others have previously shown that this strain has high levels of anxiety and low fear discrimination (8, 38). We tested the hypothesis that salient safety conditioning would reduce behavioral signs of anxiety and improve fear inhibition during novel CS− acquisition, thereby improving discrimination learning in high-anxiety animals, providing a model for future research in anxious human populations. We systematically varied the level of cue salience, given the relative difficulty in learning feature-negative cues, to directly test whether varying the conditions of learning impacted the magnitude of effects.

## 2. Materials and Methods

### 2.1 Subjects

All procedures were approved by the Hunter College and City University of New York Animal Care and Use Committee. A total of 52 129SvEv male mice (29-40g, 9-12 weeks old, Taconic BioSciences, Hudson, NY) were used in 4 experiments in this study. After arrival at the facility, animals were group housed (4/cage) for a week to acclimate, and then single housed for 1 week prior to the start of testing, when they underwent handling for 2 days (5 min/day). All animals had *ad libitum* access to food and water throughout the testing period, and were maintained on a 12 hour light/dark cycle. All testing was conducted during the light phase of the cycle.

### 2.2 Behavioral Chambers

#### 2. 2. 1 Conditioning and In-Context Retrieval Apparatus

All conditioning was conducted in a dimly lit (30 Lux) conditioning chamber (29.5 cm length x 24.8 cm width x 18.7 cm height, MedAssociates, St. Albans, VT). A house light (ENV-215M 28V, 100mA, MedAssociates) was placed in the center of the chamber ceiling (18.7 cm above floor). Depending on the group, the house light was turned on for 1 s either at the beginning or at the end of a conditioned stimulus, increasing the illumination in the chamber by 50 Lux to 80 Lux. Auditory cues were delivered via an audio speaker (ENV-224AM) located in the wall of the chamber, 10.4 cm above the floor. Shocks were delivered via a constant current aversive stimulator (ENV-414S). Side video cameras recorded the behavior for subsequent offline scoring.

#### 2. 2. 2 Innate Anxiety Chamber

To test innate anxiety and exploratory behavior, animals were placed in a grey, wooden, square enclosure (70 Lux, 50 cm sides, 50 cm wall height). The bottom of the enclosure was open and the floor was covered with grey matte paper, which was changed between subjects. Auditory cues were presented via an audio speaker (as in Section 2.2.1), placed 1m above the center of the field.

#### 2. 2. 3 Memory Retrieval Chamber

To test memory retrieval, animals were placed in a grey, rectangular enclosure (70 Lux, 45cm long × 13cm wide, 20 cm wall height). Conditioned auditory cues were presented via an audio speaker (as in Section 2.2.1 and Section 2.2.2), placed 20 cm above the floor.

### 2.3 Training and Testing Procedures

#### 2. 3. 1 Habituation (2 sessions)

One day after the second handling session (as in Section 2.2.1), each animal was brought down from the animal facility to get acclimated to the laboratory environment for 1-2 hours before being placed in the conditioning apparatus. In the apparatus, after 120 sec, the first trial began. Each subject in the Fear Conditioned and in the Safety Conditioned groups (n=13/group) received five neutral cues (4 kHz, 50ms tone pips presented at 1 Hz for 30 seconds, Pseudorandom ITIs: 60, 80, 100, or 120 s). Each subject in the Salient Beginning Safety, and Salient End Safety group (n=13/group) received five 4-kHz tones paired with a 1s house light at the beginning or the end of the tone, respectively.

#### 2. 3. 2 Phase I: Fear Conditioning Training and Testing

One day after the second habituation session, each animal returned to the conditioning apparatus. In the apparatus, after 120 sec, the first trial began. Each subject (n=13/group) received 5 cues that were each co-terminated with a shock (1 s, 0.6mA). One day after the second training session, mice were placed back in the conditioning apparatus without any shock delivery for testing. As during the conditioning sessions, the first non-reinforced tone trial was presented after 120sec in the Conditioning apparatus (2.2.1). Fear responses in the Conditioning apparatus were quantified by measuring the amount of time spent freezing during the 30 s prior to each CS onset (contextual fear) and during the 30 s during CS presentations. Complete locomotor cessation, for at least 1s, was counted as freezing.

##### Phase I: Safety Conditioning Training and Testing: Safety, Salient End Safety, Salient Beginning Safety

One day after the second habituation session, each animal was placed in the conditioning apparatus. After 120 sec in the Conditioning apparatus (2.2.1), the first trial began, and each animal (n=13/group) received 5 tone cues that were explicitly unpaired from 5 shock US presentations (1s, 0.6mA, mean interval between US and all proximal CSs, 49s; range, 40 to 60 sec). Shock delivery never occurred during the 30-second tone, but shock output could occur during the ITI. However, the offset of the tone did not predict shock onset as the shocks could be delivered anywhere from 0-2 times on any given ITI. In addition to the tone CS, mice in the Salient End Safety and Salient Beginning Safety conditions also received a house light output co-administered for 1 second either with each tone offset (Salient End group, n=13) or tone onset (Salient Beginning group, n=13). We reasoned that light paired with the onset of the safety tone would add salience to the subsequent safe period, whereas lights presented at the offset of the safety tone would add salience to the subsequent period during which a shock could occur. One day after the second training session, mice were placed back in the Conditioning apparatus and were presented with light-tone cues without any shock delivery. Fear responses in the conditioning chamber were quantified by measuring the amount of time spent freezing during the 30 sec prior to CS onset (contextual fear), and during the 30 sec of CS presentations. Complete locomotor cessation, for at least 1sec, was counted as freezing.

#### 2. 3. 3 Phase I: Open Field Exploration in the presence of fear or safety conditioned cues

One day after the testing session, each animal was placed in the open field (2.2.2) for a total of 6 minutes. The mice explored the open field with alternating 1-minute periods of no CS, followed by 1-minute of CS presentations (3 alterations, 60 seconds each).

#### 2. 3. 4 Phase II: Differential Fear Conditioning Training and Testing

Three days after the open field exposure, mice were differentially fear conditioned on new cues for three consecutive days (1 session per day). Each day, twelve auditory trials (6CS+ and 6CS−: 1kHz or 7kHz 50-ms tone pips occurring at 1Hz, 30 sec counterbalanced) were pseudorandomly presented (ITI, 60–120 s). Each CS+ co-terminated with a 0.6 mA, 1 sec footshock US, whereas no footshock was paired with the CS−. On day 4, mice were placed in the memory retrieval chamber (Section 2.2.3), and presented with twelve pseudorandomly distributed CS+ and CS− trials without shock delivery.

### 2.4 Data Analyses

Our experimental groups to directly compare Fear and Safety conditioning on subsequent behavior consisted of 13 male mice per group (Phase I training). Cohorts of 3 – 5 *Fear conditioned* mice were trained alongside cohorts of 3 – 5 *Safety, Salient End Safety,* and *Salient Beginning Safety* mice for a total of 3 replications. Custom-made Matlab (Natick, MA) scripts were used to cut out tone and pre-tone periods from the video recordings, and the identities of the animals were scrambled. The tone and pre-tone videos were then assessed manually offline by a coder blinded to training conditions for periods of freezing, defined as complete immobility lasting at least 1 second, as a measure of innate defensive behavior (39). The total time spent freezing during each 30s cue as well as during the 30s preceding the cue was quantified and expressed as a percentage.

To determine the effects of Phase I training on exploratory behavior in an anxiogenic environment, we recorded animal behavior in the open field (2.2.2). Measuring the total time animals spent moving inside the center or periphery of the open field in six 1-minute intervals. We assessed the percent time spent in the center of the field, and the distance that the animals moved. The following criteria for determining center and periphery were used: 0-10cm from the wall was considered periphery, 10-50cm from the wall of the open-field was considered the center. Percent time in the center or the periphery and distance traveled were computed offline, using AnyMaze software (Wood Dale, IL) with mapped dimensions of the field using videos recorded from an overhead camera. Alternating 60-second intervals of no-tone and tone were repeated three times for a total of six minutes. The final 2 min were excluded from analysis due to animal inactivity.

To determine the effects of training condition on subsequent discrimination learning, freezing behavior was measured in the memory retrieval chamber (2.2.3) during periods of CS+ and CS− tone presentation. The tone and pre-tone periods were cut using a custom-written Matlab script, and the animals’ identities were scrambled. Fear behavior was assessed manually offline by a coder blind to training condition from the videos, by measuring the total time spent freezing during each 30s cue, expressed as a percentage. A discrimination score was quantified for each mouse based on their averaged freezing response to 6 CS+ and 6 CS− trials. The discrimination score was computed as %Freezing CS+ −% Freezing CS−, and also normalized to total freezing, by computing % Freezing CS+ -% Freezing CS−/ Total %Freezing. Behavioral data were analyzed with one-way or two-way repeated measures ANOVAs, with training condition as the independent factor, and trial number as the repeated factor, followed by *post hoc* Sidak’s, Tukey’s or Dunnett’s multiple comparisons tests with GraphPad Prism 8 and verified on SPSS. *P−*values were adjusted for multiple comparisons.

## 3. Results

### 3.1 Salient Beginning Safety conditioning is the most effective training for in-context fear suppression

All mice underwent habituation (2 days), and then were randomly placed in one of four groups: 1. Fear conditioning where the shock US co-terminated with tone CS; 2. Safety conditioning where the tone CS and shock US were explicitly unpaired; 3. Salient Beginning Safety conditioning where tone CS and shock US were explicitly unpaired and a light CS was co-presented for 1 s at the beginning of tone CS; 4. Salient End safety conditioning where the tone CS and shock US were explicitly unpaired and a light CS was co-presented for 1s at the end of tone CS (Section 2.3.2, Fig. 1a). All groups showed increased tone-evoked freezing by day 2 of training (Figure 1b). A two-way repeated measures ANOVA on trial freezing by group showed no significant interactions (*F*(12,176) = 1.167, *p* = 0.31), indicating that animals in all groups learned the combined aversive nature of the cues and context during training.

**Figure 1:**
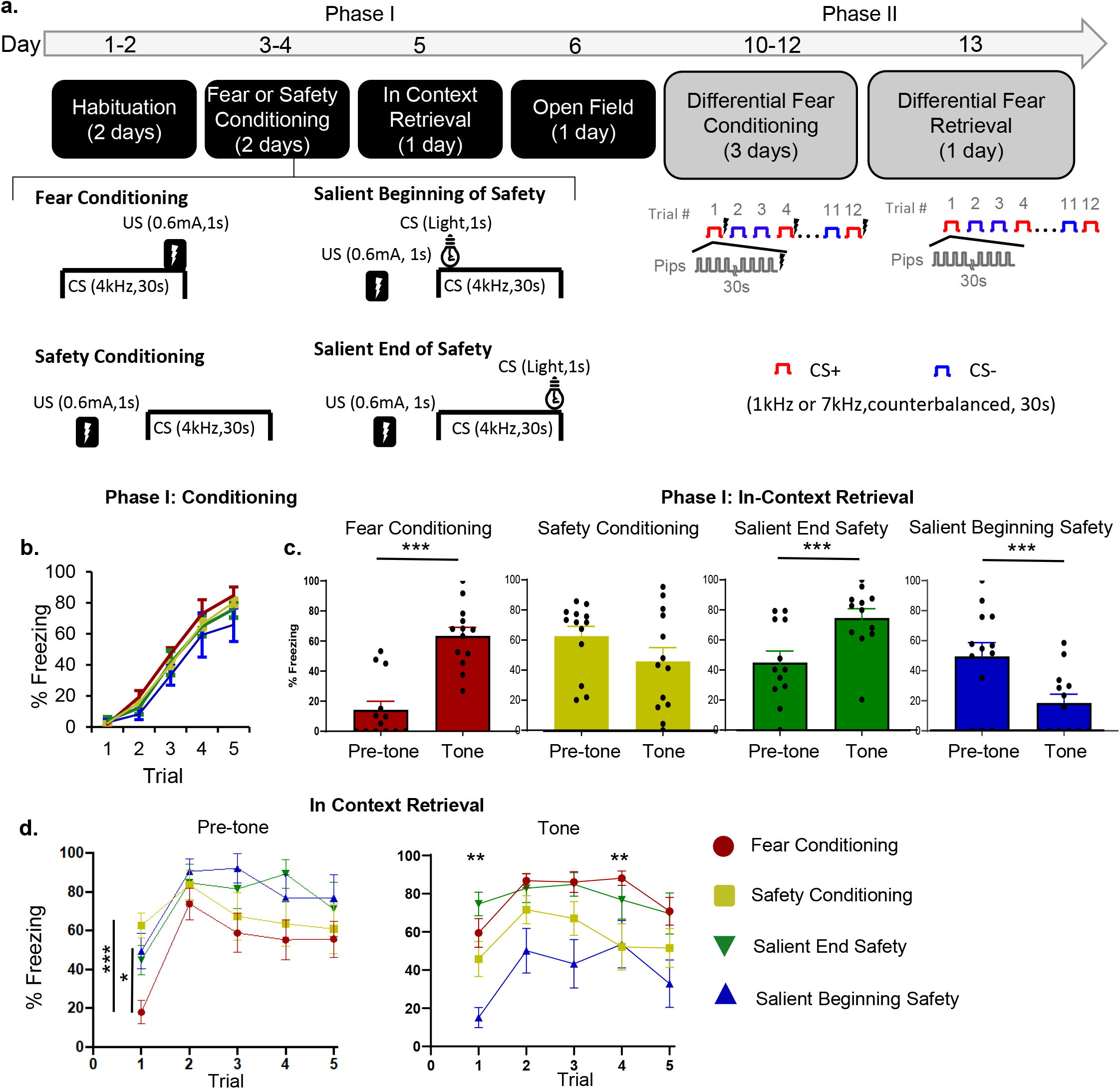
Salient Beginning Safety Conditioning is the most effective safety training for in context fear inhibition. **(a)** Timeline of behavioral testing. Mice were first habituated to the conditioning context for 2 days (Days 1-2), and then either fear or safety conditioned (5 trials per day, Days 3-4). Fear conditioning consisted of CS1 (50 ms long 4kHz tone-pips, once a second, lasting 30 sec) co-terminating with a shock US (0.6mA, 1 s). Safety conditioning consisted of CS1 presentations that were explicitly unpaired with the US (mean interval between US and all proximal CSs, 49 sec; range, 40 to 60 sec). The Salient Beginning Safety Conditioning was structured in the same way as the Safety conditioning, the only difference being a CS2 (houselight) that was co-presented for 1 sec at the beginning of CS1. The Salient End Safety Conditioning group was presented with CS2 co-terminating with the end of CS1 for 1 sec. The next day, mice were tested for in-context fear retrieval (Day 5), and on Day 6, mice were placed in the Open Field to test for the effects of fear or safety conditioning in Phase I on exploration of a novel anxiogenic context. After a 3 day break (Day 7-9), mice underwent differential fear conditioning for 3 days (12 trials per day, Days 10-12), and then were placed in a new context for fear discrimination retrieval (Day 13). (**b**) Defensive freezing for all groups during conditioning (Day 3). (**c**) In-context retrieval of the learned associations. Pre-tone freezing for all groups in the 30s preceding the first tone and during the 30s of the first tone. (**d**) In-context retrieval behavior across all trials. Left, defensive freezing for all groups during the 30s pre-tone period prior to each tone. Right, defensive freezing for all groups on each trial. ***, p<0.0001; **, p<0.001, *p<0.05.

To assess whether animals learned cue-specific fear and safety associations, we tested in-context retrieval of the tone CS the day after training (Fig. 1c). A two-way ANOVA comparing group by tone-period showed a main effects of group (*F*(3, 48) = 4.176, *p* = 0.01) and tone-period (*F*(1, 48) = 4.851, *p* = 0.03), with a significant interaction between pre-tone and tone in the fear conditioned group where the tone significantly increased defensive freezing (Fear group pre-tone freezing = 14.33±5.62%, Fear group tone freezing = 63.44±5.72%, *p* < 0.0001). The Safety group that didn’t have a salience component during training did not effectively inhibit contextual pre-tone freezing (Safety group pre-tone freezing = 62.65±6.47%, Safety group tone freezing = 45.79±9.13%, *p* = 0.077), suggesting that simple safety training is not sufficient to decrease fear in high anxiety mice. On the contrary, the Salient Beginning safety group showed a significant decrease in contextual freezing during the tone (Fig. 1c, Salient Beginning safety group pre-tone freezing = 49.56±9.14%, Salient Beginning safety group tone freezing = 18.56±5.78%, *p* = 0.0002). Notably, there was no significant difference in pre-tone freezing between the Safety and the Salient Beginning safety groups (Safety pre-tone freezing = 62.65±6.47%, Salient Beginning pre-tone freezing=49.56±9.14%, *p* = 0.71). However, the Salient Beginning safety group froze significantly less during the tone than the Safety group (Salient Beginning group tone freezing = 18.56±5.78%, Safety group tone freezing = 45.79±9.13%, *p* = 0.04). Animals in the Salient End safety group behaved similarly to the Fear Conditioned animals, whereby the cue evoked more freezing than context alone, although this effect almost reached significance (Fig. 1c, Salient End safety pre-tone freezing = 45.21±7.03%, Salient End safety tone freezing = 74.78±5.65%, *F*(1,12) = 4.42, *p* = 0.057). Thus, our data demonstrate that for high anxiety animals, Salient Beginning Safety training was the most effective paradigm for inhibiting fear in the training context.

Overall, there was a significant interaction of pre-tone 1(contextual) freezing by group (*F*(1,12)=18.14, *p*=0.0011, pre-tone freezing in the Fear group = 14.33±5.62%, the Safety group = 62.65±6.47%, the Salient Beginning safety group = 49.56±9.14%, the Salient End safety group = 45.21±7.03%), demonstrating that prior to trial 1, fear conditioned animals froze less to the context than safety conditioned animals in all three groups (Figure 1c,d).

A repeated measures ANOVA for pre-tone freezing by group across trials showed that there was no main effect of group (*F*(3,40) = 2.591, *p* = 0.07), but there was a main effect of trial (*F*(3.52,140.9) = 13.88, *p* < 0.0001), and there was a trial by group by interaction, with significantly less pre-tone 1 freezing in the Fear Conditioned group than in the Safety group (*p* < 0.0001), and in the Fear Conditioned group than the Salient Beginning safety group (*p* = 0.041). After pre-tone 1, ITI freezing in fear conditioned animals increased and was not different from that of safety trained groups (Figure 1d), underscoring the anxious phenotype of this strain.

However, animals in the Salient Beginning safety group showed better fear suppression than other safety groups during tone trials across the session, in addition to trial 1 described above (Fig. 1c). A repeated measures ANOVA for trial freezing by group showed that there was a main effect of trial (*F*(3.69,173.4) = 8.49, *p* < 0.0001), and a main effect of training (*F*(3,47) = 8.47, *p* = 0.0001), but there was no interaction (*F*(4,188) = 0.42, *p* = 0.79). *Post-hoc* Tukey’s multiple comparisons testing revealed that across 5 trials (Fig. 1d), the Fear Conditioned group froze equally high to the tone as the Safety conditioned group, and the Salient End safety conditioned group on all trials. However, the Salient Beginning safety group froze significantly less during tone 1 (*p* = 0.0086), and tone 4 (*p* = 0.04) than the Fear Conditioned groups (Fig. 1d). These findings indicate that the Salient Beginning safety group showed tone-evoked fear suppression throughout the testing period.

### 3.2 Safety training improves exploration of an anxiogenic environment

Next, we wanted to investigate whether the conditioned safety and fear cues modulate behavior in a novel, anxiogenic environment. To this end, animals were placed in the open field (Day 6), where they spent 6 minutes, alternating between one-minute trials with no cue present and with the trained 4kHz cue presented via a speaker (Section 2.3.3, Fig. 1a). Previous work has shown that open field exploration decreases during short exposures to the environment (but notably increases after longer exposures) (40), and similarly in our experiment total distance traveled in all groups decreased to less than 1m per trial after minute 4 (data not shown). Given that we were interested in how the cue modulates behavior rather than long-term exploration of the field (40), we analyzed behavior during the first two CS Off and CS On periods (Figure 2a-b). Overall, the Fear Conditioned cue modulated behavior more than the safety cues, regardless of safety training protocol. A two-way ANOVA comparing tone presentation x training group demonstrated a main effect of tone on distance traveled in the open field (*F*(1,96)=68.20, *p* < 0.0001) without a main effect of training condition (*F*(3,96) = 0.071, *p* = 0.98), but with a significant interaction (*F*(3,96) = 6.18, *p* = 0.0007, Fig. 2b-c). *Post-hoc* Tukey’s multiple comparisons tests showed that animals in the Fear Conditioned group moved more on the open field when the cue was off (*p* < 0.0001), whereas animals in the safety trained groups did not modulate movement with the tone (Figure 2c, *p*>0.05). Likewise, a one-way ANOVA comparing the distance traveled through the anxiogenic center of the open field across the four conditioning groups showed a significant effect of conditioning (*F*(3,47) = 7.67, *p* < 0.001). A *post-hoc* Tukey’s multiple comparisons test showed that the Fear Conditioned animals moved significantly more in the center open of the field during the Tone Off than the Tone On period, whereas animals in all the safety trained groups did not alter their movement in the center of the open field based on tone presence (Fig. 2d, *p* < 0.01).

**Figure 2.**
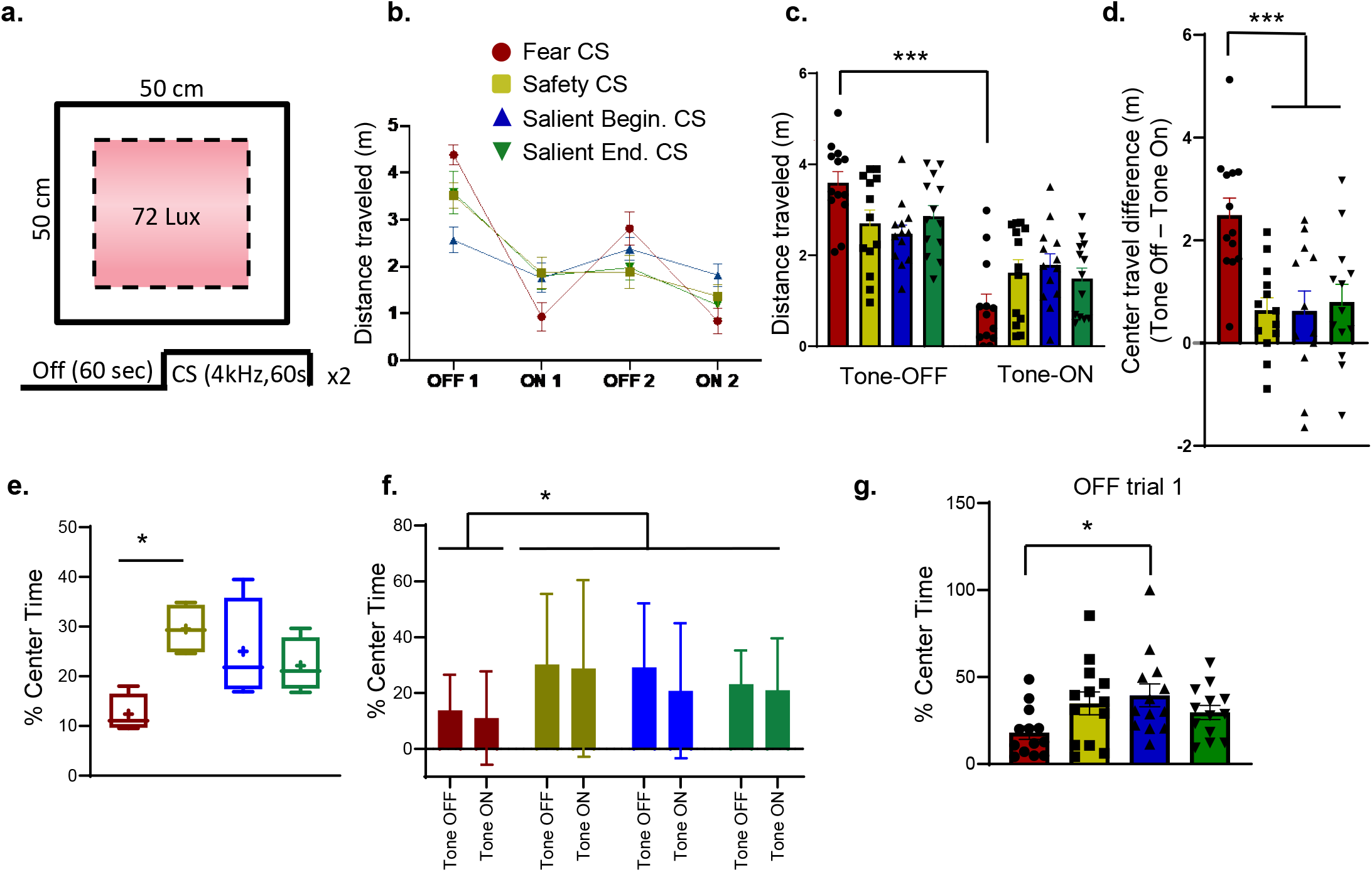
Safety training improves exploration of a novel anxiogenic environment. (**a**) Protocol and schematic of the open-field used for innate anxiety testing. (**b**) Exploratory behavior measured as distance traveled from all groups throughout the entire field (meters). (**c**) Average distance traveled by each group throughout the field during Tone-off and Tone-on periods. In the Fear Conditioned group, the average travel distance during Tone-on is significantly lower (0.88m±0.26) than during the Tone-off periods (3.60m±0.24). (**d**) Change in travel distance within the anxiogenic center of the field. Change measured as center distance travelled during Tone-Off – Tone-On period. The average difference is significantly lower in the safety groups; Safety = 0.64m ±0.24, Salient Beginning safety = 0.63m ±0.36, Salient end safety = 0.80m ±0.34 than the Fear Conditioned group 2.49m ±0.33. (**e**) Box and whisker plot showing percent center time for each group during Tone-on periods. (**f**) Mean percent center time during Tone-off and Tone-on trials. (**g**) Mean percent center time during Trial-Off 1 (the first 60 seconds in the open field). Percent time in the center is significantly higher in the Salient Beginning Safety Group (39.49±6.58%) compared to the Fear Conditioned Group (18.04±3.83%).

We then compared whether fear or safety conditioned cues modulate the percent of time animals spent in the anxiogenic center of the open field. A one-way ANOVA comparing the overall percent time spent in the center x training group showed a significant main effect of training (*F*(3,12) = 4.76, *p* = 0.02, Fig. 2e-f). A *post-hoc* Tukey’s multiple comparisons test showed that the fear conditioned animals spent less time in the center overall than the safety conditioned group (*p* = 0.015, Fig. 2e). Further, a two-way ANOVA comparing center time exploration as modulated by cue in each conditioning group, showed that there was a significant effect of training (*F*(3,96) = 2.96, *p* = 0.03), but no effect of tone (*F*(1,96) = 0.76, *p* = 0.38), and no interaction (*F*(3,96) = 0.14, *p* = 0.93), suggesting that the tone did not modulate animals’ exploration of the center but previous training did. Again, p*ost-hoc* Tukey’s multiple comparisons showed that overall the Fear Conditioned animals spent less time in the center than the safety conditioned groups (Fig. 2e, *p* = 0.029).

Given that the effects of exploration were independent of tone, but rather depended on the type of conditioning that animals underwent in Phase I training, we analyzed exploration of the anxiogenic center independent of tone. We used a one-way ANOVA to evaluate the percent center time during the first minute of open field exposure across groups, prior to animals hearing tones. This analysis revealed a significant difference between groups (*F*(3,48) = 2.9, *p* = 0.04), suggesting that previous training affected center exploration time. *Post-hoc* Tukeys multiple comparisons tests revealed that the Fear Conditioned group explored the center significantly less during the first minute of exposure (Fig. 2g, 18.04% ± 13.8) than the Salient Beginning safety conditioned group (39.49%±23.8%), p=0.035). These findings indicate the Salient Beginning safety training improves subsequent exploration of an anxiogenic environment in a high anxiety strain. However, presentation of the safety cue does not increase exploration or movement in a new anxiogenic environment.

#### 3.3 Salient Beginning Safety trained mice discriminate the CS− better on new differential fear conditioning task

To assess whether exposure to fear or safety conditioning affects subsequent fear discrimination learning, we implemented a Phase II training where we differentially fear conditioned animals from all four Phase I training groups to new cues - 1 and 7kHz tones, that were counterbalanced as the CS+ and CS− (Fig. 1a, Days 10-12). Animals were then placed in a new memory retrieval apparatus (Section 2.2.3), and presented the CS+ and CS− (Fig. 1a, Day 13). A two-way ANOVA measuring cue-evoked defensive freezing x Phase I training showed a significant main effect of cue (*F*(1,96) = 43.96, *p* < 0.0001), and Phase I training (*F*(3,96) = 4.62, *p* = 0.005), as well as an interaction between them (*F*(3,96) = 6.72, *p* = 0.0004). *Post-hoc* Tukey’s multiple comparisons tests showed that animals who went through Salient Beginning Safety conditioning in Phase I were the only group to have significantly different defensive freezing to the CS+ and the CS− in Phase II (Fig. 3a, *p* < 0.0001). All the other groups showed similar average freezing to the two tones (Fig. 3a, Fear Conditioning group, *p* = 0.98; Safety group, *p* = 0.22; Salient End safety group, *p* = 0.07).

**Figure 3:**
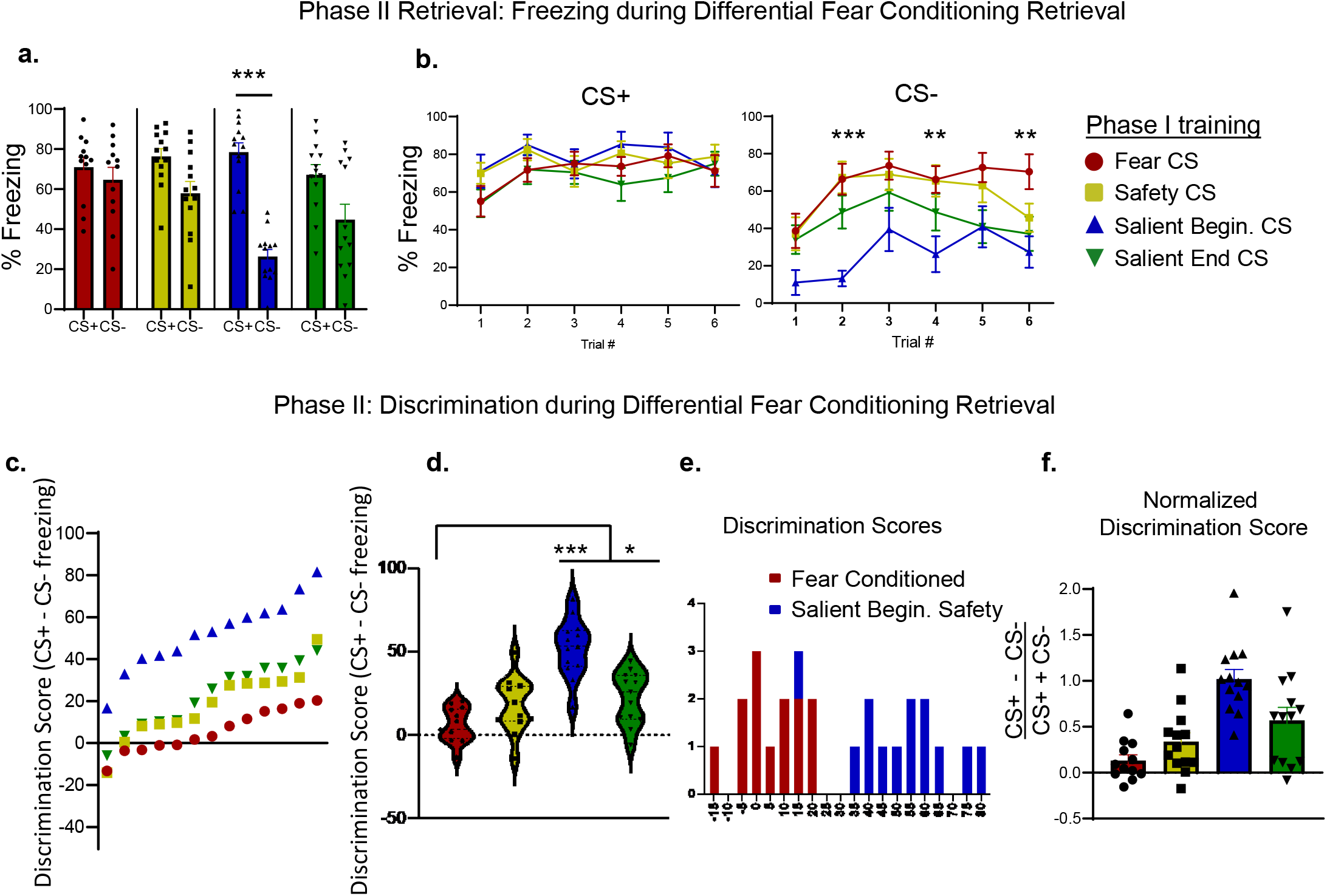
Salient Beginning Safety conditioned mice discriminate well during novel differential fear retrieval. (**a**) Percent CS+ (Left panel) and CS− (Right panel) freezing on each trial of a fear discrimination retrieval task for animals that in Phase I were either Fear conditioned (red) or Safety Conditioned in one of three groups (Safety– yellow, Salient Beginning safety– blue, Salient End safety– green). (**b**) Average trial freezing during differential fear conditioning retrieval in all groups (**c**) Discrimination scores, quantified as percent freezing to the CS− subtracted from percent freezing to the CS+, shown for all subjects (each marker shows individual discrimination score for each animal, color coded by Phase I training). (**d**) Violin plot showing discrimination score distributions across all four groups of animals. The median, 25^th^ and 75^th^ quartile, and the minimum and maximum are shown for each group. (**e**)Histogram showing the distribution of Phase II discrimination scores in animals that were either Fear Conditioned (red) or Salient Beginning Safety Conditioned (blue) in Phase I. The average discrimination score in the fear conditioned group is significantly lower (5.62±2.85%) than in the Salient Beginning safety conditioned group (52.10 ±4.79%). (**f**) Discrimination scores normalized for total freezing in all groups. The Salient Beginning Safety and Salient End Safety conditioned groups show significantly higher normalized discrimination than the fear conditioned group (Fear Conditioned in Phase I = 0.13±0.06, Safety conditioned in Phase I = 0.34±0.10, Salient Beginning safety conditioned in Phase I = 1.02.56±0.11, Salient End safety conditioned in Phase I = 0.57±0.14. mean ± SEM). The Salient Beginning Safety conditioned groups shows the highest discrimination.

A repeated measures ANOVA comparing defensive freezing across CS+ trials x Phase I training showed a main effect of tone number (*F*(4.57,215) = 3.44, *p* = 0.007), but no main effect of group (*F*(3,47) =1.22, *p* = 0.31), and no interaction (*F*(15,235) = 1.24, *p* = 0.29). Thus, regardless of training in Phase I, all groups showed equivalent levels of freezing during CS+ retrieval in Phase II (Fig. 3b). However, when comparing levels of CS− evoked freezing across groups, a repeated measures ANOVA showed that there was a main effect of tone number (*F*(4.49,210.9) = 8.86, *p* < 0.0001), and group (*F*(3,47) = 7.79, *p* = 0.003). *Post-hoc* comparisons using Tukey’s multiple comparisons test showed that on retrieval trial 2 (*p* < 0.001), trial 4 (*p* = 0.015), and trial 6 (*p* = 0.012) animals that were fear conditioned in Phase I froze more to the CS− than those that were Salient Beginning Safety conditioned in Phase I (Fig. 3b). Thus, animals that were safety conditioned using the Salient Beginning protocol during Phase I, showed the lowest freezing during the CS− retrieval trials, while continuing to freeze to the CS+ at equally high levels as the other groups (Fig. 3a-b).

We used a Discrimination Score (percent freezing to the CS+ minus percent freezing to the CS−) to capture the difference in the percent freezing to the two conditioned cues. A one-way ANOVA showed significant differences in discrimination scores among the four Phase I training groups (*F*(3,48) = 21.9, *p* < 0.0001, Fig. 3c-d). *Post-hoc* analyses showed that the group that was Fear Conditioned in Phase I, discriminated significantly less than the group that was Salient Beginning safety conditioned in Phase I (*p* < 0.0001), as well as Salient End safety conditioned in Phase I (*p* = 0.03). However, the previously Fear Conditioned and regular Safety conditioned groups did not discriminate differently (*p* = 0.15). Notably, the Salient Beginning safety conditioned group had a significantly higher discrimination score (52.1±4.8) than the Fear Conditioned group (5.6±2.9, t-test, *p* < 0.0001), with a distribution of discrimination scores that only slightly overlapped with the fear conditioned group (Figure 3e).

Next, we normalized the discrimination score by overall freezing during retrieval (CS+ − CS−/ CS+ + CS−), in order to account for any differences in total freezing between groups. A one-way ANOVA comparing normalized discrimination scores showed significant differences in normalized discrimination between groups (Fig. 3f, *F*(3,48) = 13.24, *p* < 0.0001). *Post-hoc* Tukey’s multiple comparisons tests showed that, similarly to the non-normalized case, the group that was Fear Conditioned in Phase I, discriminated significantly worse than the group that was Salient Beginning safety conditioned in Phase I (Fig. 3f, *p* < 0.0001). Likewise, the Fear Conditioned group discriminated worse than the Salient End Safety conditioned group (*p* = 0.022), but there was no difference between the Fear Conditioned and the regular Safety conditioned groups (*p* = 0.49). These findings suggest that high anxiety animals benefit from undergoing Salient Beginning safety training in order to better discriminate the CS− during subsequent learning. We find that boosting the salience of the safety cue with another cue (here, a brief light) at the beginning but not the end of the safety cue improved novel discrimination in this anxious strain of mice.

## 4. Discussion

The aim of this study was to test whether safety learning improves discrimination learning in 129SvEv male mice, a strain that typically shows high anxiety and low fear discrimination. Here, we compared how safety cues trained in one of three ways modulate subsequent exploration and learning: (i) the safety cue that is explicitly unpaired with the occurrence of the shock, (ii) a safety cue that is unpaired with the shock but is made more salient by the co-occurrence of a 1-sec light at cue onset, and (iii) a safety cue that is unpaired with the shock but cue termination is made more salient with 1-second light co-occurring at cue offset. Notably, all groups were exposed to an equal number of shocks throughout training, thereby equalizing the physical stress that animals underwent. Despite the equal stress exposure, our findings demonstrate that in the 129SvEv strain, exposure to salient safety cues (Fig. 1) improves subsequent fear discrimination learning by decreasing the defensive freezing to a new, non-threatening CS− (Fig. 3). Notably, male mice were used for the most direct comparisons to our previous discrimination learning data with this strain (8, 35). However, anxiety disorders are more prevalent in women than in men (41–43), emphasizing the need to repeat this experiment in female mice, as the effects of safety training on discrimination learning needs to be evaluated in both sexes.

Our results show that using a non-salient safety stimulus during initial safety training is not an effective suppressor of fear in the training context for this high anxiety strain of mice (Fig. 1). This is in contrast to the same protocol leading to successful downregulation of fear by the non-salient safety cue previously shown in C57Bl/6 mice (14), who present with a less anxious phenotype. Given that the 129SvEv mice show more behavioral similarities to high anxiety clinical populations, we wanted to develop safety training paradigms that downregulate defensive behavior in the training context more effectively. Increasing the salience of the safety signal at the beginning of the cue, likely driving more attention to the safety cue, proved to be the most effective means for making the safety cue a potent inhibitor or fear in this high anxiety strain. In contrast, increasing the salience of the safety cue at the end, highlighted the onset of the uncertainty associated with the shock period, and instead of downregulating defensive behavior, this cue increased it (Fig. 1c). Thus, manipulating attention at different points in the safety cue has drastically different effects on how it regulates fear suppression and expression in high anxiety animals.

The 129SvEv strain used here shows similar patterns of fear generalization (8) to anxious humans, in that heightened anxiety disrupts effective learning about the CS− during discrimination (4–6). Our findings suggest that enhanced safety training, potentially by increasing attention to the onset of a cue signaling safety, improves subsequent discrimination of other non-threatening cues, whereas the non-salient safety training did not significantly improve discrimination (Fig. 1a, Fig. 3). Notably, our results show that, when animals retrieve safety cues where the end of the cue was paired to a brief light signaling the end of safety, defensive freezing is high (Fig. 1c-d), likely because anxiety is amplified by increased attention to the end of the safety period, and the onset of uncertainty regarding the delivery of an aversive stimulus. In humans, intolerance of uncertainty is an overemphasized feeling of discomfort with uncertainty that something aversive may happen, and has been associated with heightened anxiety (44–46) Similarly, our results showed that mice that underwent Salient End Safety conditioning, may have experienced increased anxiety about the safety cue because when played back in the same context, it highlighted the uncertain, anxiety-inducing period of shock occurrence and increased defensive freezing, as for the fear conditioned animals (Fig. 1). However, even though the Salient Beginning Safety trained group showed the most robust learning during discrimination learning (Phase II), the Salient End Safety group still showed better overall discrimination than the initial Fear Conditioned group (Fig. 3). This finding suggests that modulation of attention to the beginning versus the end of safety cues can have lasting clinical implications for anxiety.

Discrimination learning depends on multiple stages of stimulus processing, and stress-or fear-induced disruption of discrimination can also occur at several relay points of neural computation (47). For example, at the sensory level, recordings in the auditory cortex show that fear conditioning broadens cortical receptive fields for the aversive cue, and increases fear generalization (7, 48). Underscoring the importance of inhibitory cells in sculpting auditory cortex output during fear conditioning, the aversive cue drives local inhibitory microcircuits that disinhibit pyramidal cell activity (49). Furthermore, inhibiting parvalbumin expressing interneurons of the auditory cortex impairs frequency selectivity in auditory pyramidal cells, and enhances behavioral fear generalization (50). It is not yet clear how safety training affects sensory processing or how inhibitory neurons shape auditory discrimination during safety, however this is one potential route for safety training to affect behavioral output.

Another important aspect to note about anxiety in relation to aberrant attentional processing is that anxiety is associated with exaggerated, sustained attention to threat (51–54), termed attention bias (54, 55). Given that basal forebrain cholinergic activity contributes to attention and memory (56–59), increased attention bias could disrupt cholinergic function, contributing to stimulus generalization. In keeping with this, disinhibition observed in the auditory cortex during the aversive stimulus depends on cholinergic inputs from the basal forebrain (49, 60). Accordingly, cholinergic neurons respond to cues during training, and reshape receptive fields in the auditory cortex, even during trace conditioning when the CS and US are seconds apart (48, 60). Likewise, low levels of stimulation of the cholinergic nucleus basalis decreases the specificity of the cue-evoked defensive response, whereas higher stimulation levels make the response more cue-specific, and increase gamma frequency in the auditory cortex (48). Our data suggest that to reap the benefits of safety learning, and to counteract attention bias in a high-anxiety mouse strain, the onset of the safety cue should be amplified in order to increase attention to the safety period. It’s possible that increased attention to the safety cue also modulates cholinergic input to the auditory cortex, shaping stimulus generalization curves.

The mPFC, and BLA are two more regions that encode safety cues, and have a prominent role in fear discrimination learning (15, 22, 31, 34, 37, 61–65). There are separate BLA neurons that encode the CS+ and the CS− during discrimination learning, as well as separate neurons that encode the CS+ and the safety cue (22, 23, 26, 64), suggesting that neurons encoding the CS− and the safety cue may rely on at least partially overlapping circuits. In the case of a discriminated CS−, there is increased communication between the mPFC and the BLA, when mPFC theta oscillations entrain BLA firing, and increase mPFC coupling with BLA gamma oscillations (8, 35), a communication pattern that develops through discrimination learning (34, 65). It is not known whether the same pattern of prefrontal communication with the amygdala develops during explicit safety cue learning as during CS− discrimination. Our finding that safety training improves CS− discrimination in subsequent learning suggests that similar patterns of communication in this circuit may be active. It is possible that after engaging mPFC-BLA circuits during initial safety training, there is increased plasticity during subsequent CS− acquisition.

Previous work demonstrated that in the C57Bl/6 strain of mice, safety cues modulate subsequent behavior, for example increasing center exploration in an open-field, and motivating animals in a place preference task (15). We also demonstrate that safety trained groups, regardless of training protocol, show increased exploration in an open field compared to fear conditioned animals. However, cue presentation only modulated behavior in the fear conditioned group, whereas changes in exploration were independent of cue for the safety trained groups, suggesting that safety training affected innately driven exploration. Our open field exposure was short-lasting (6 minutes) because we were interested in identifying how learned fear-and safety-cues modulate exploration. However, previous work has shown that during longer periods of open field exposure, rodents demonstrate complex exploration and foraging patterns (66). Given our finding that safety trained groups showed increased initial exploration relatively to the fear trained group, it would be interesting to know whether safety training improves the exploration and foraging success with a longer exposure to the open field.

In sum, we demonstrate that safety training has positive consequences on subsequent fear discrimination learning, and on exploration in a novel, anxiogenic environment. Increasing the salience of the safety cue by adding a second cue at its onset improved safety conditioning, and improved performance on subsequent discrimination learning, suggesting that boosting attention to safety cues for high anxiety individuals can be clinically beneficial.

## Acknowledgements

We thank Sandra Talbot for her help with this project.

## References

1. Aldao A, Nolen-Hoeksema S, Schweizer S. Emotion-regulation strategies across psychopathology: A meta-analytic review. Clin Psychol Rev. 2010;30(2):217–37.

2. Borkovec TD, Alcaine OM, Behar E. Avoidance Theory of Worry and Generalized Anxiety Disorder. Generalized anxiety disorder: Advances in research and practice. New York, NY, US: Guilford Press; 2004. p. 77–108.

3. Duits P, Cath DC, Lissek S, Hox JJ, Hamm AO, Engelhard IM, et al. Updated meta-analysis of classical fear conditioning in the anxiety disorders. Depress Anxiety. 2015;32(4):239–53.

4. Lissek S, Kaczkurkin AN, Rabin S, Geraci M, Pine DS, Grillon C. Generalized anxiety disorder is associated with overgeneralization of classically conditioned fear. Biol Psychiatry. 2014;75(11):909–15.

5. Dymond S, Dunsmoor JE, Vervliet B, Roche B, Hermans D. Fear Generalization in Humans: Systematic Review and Implications for Anxiety Disorder Research. Behav Ther. 2015;46(5):561–82.

6. Lissek S, Powers AS, McClure EB, Phelps EA, Woldehawariat G, Grillon C, et al. Classical fear conditioning in the anxiety disorders: a meta-analysis. Behav Res Ther. 2005;43(11):1391–424.

7. Aizenberg M, Geffen MN. Bidirectional effects of aversive learning on perceptual acuity are mediated by the sensory cortex. Nat Neurosci. 2013;16(8):994–6.

8. Likhtik E, Stujenske JM, Topiwala MA, Harris AZ, Gordon JA. Prefrontal entrainment of amygdala activity signals safety in learned fear and innate anxiety. Nat Neurosci. 2014;17(1):106–13.

9. Resnik J, Sobel N, Paz R. Auditory aversive learning increases discrimination thresholds. Nat Neurosci. 2011;14(6):791–6.

10. Schechtman E, Laufer O, Paz R. Negative valence widens generalization of learning. J Neurosci. 2010;30(31):10460–4.

11. Shalev L, Paz R, Avidan G. Visual Aversive Learning Compromises Sensory Discrimination. J Neurosci. 2018;38(11):2766–79.

12. Christianson JP, Fernando AB, Kazama AM, Jovanovic T, Ostroff LE, Sangha S. Inhibition of fear by learned safety signals: a mini-symposium review. J Neurosci. 2012;32(41):14118–24.

13. Crowell CR, Bernhardt TP. The feature-positive effect and sign-tracking behavior during discrimination learning in the rat. Animal Learning & Behavior. 1979;7(3):313–7.

14. Sainsbury RS, Jenkins HM. Feature-positive effect in discrimination learning. Proceedings of the Annual Convention of the American Psychological Association. 1967;2:17–8.

15. Rogan MT, Leon KS, Perez DL, Kandel ER. Distinct neural signatures for safety and danger in the amygdala and striatum of the mouse. Neuron. 2005;46(2):309–20.

16. Sangha S, Diehl MM, Bergstrom HC, Drew MR. Know, safety, no fear. Neurosci Biobehav Rev. 2020;108:218–30.

17. Hendry DP. Conditioned inhibition of conditioned suppression. Psychonomic Science. 1967;9(5):261–2.

18. Duvarci S, Bauer EP, Pare D. The bed nucleus of the stria terminalis mediates inter-individual variations in anxiety and fear. J Neurosci. 2009;29(33):10357–61.

19. Foilb AR, Christianson JP. Serotonin 2C receptor antagonist improves fear discrimination and subsequent safety signal recall. Prog Neuropsychopharmacol Biol Psychiatry. 2016;65:78–84.

20. Marcinkiewcz CA, Mazzone CM, D’Agostino G, Halladay LR, Hardaway JA, DiBerto JF, et al. Serotonin engages an anxiety and fear-promoting circuit in the extended amygdala. Nature. 2016;537(7618):97–101.

21. Sangha S, Robinson PD, Greba Q, Davies DA, Howland JG. Alterations in, reward, fear and safety cue discrimination after inactivation of the rat prelimbic and infralimbic cortices. Neuropsychopharmacology. 2014;39(10):2405–13.

22. Sangha S, Chadick JZ, Janak PH. Safety encoding in the basal amygdala. J Neurosci. 2013;33(9):3744–51.

23. Genud-Gabai R, Klavir O, Paz R. Safety signals in the primate amygdala. J Neurosci. 2013;33(46):17986–94.

24. Herry C, Ciocchi S, Senn V, Demmou L, Muller C, Luthi A. Switching on and off fear by distinct neuronal circuits. Nature. 2008;454(7204):600–6.

25. Kim WB, Cho JH. Encoding of Discriminative Fear Memory by Input-Specific LTP in the Amygdala. Neuron. 2017;95(5):1129–46 e5.

26. Grewe BF, Grundemann J, Kitch LJ, Lecoq JA, Parker JG, Marshall JD, et al. Neural ensemble dynamics underlying a long-term associative memory. Nature. 2017;543(7647):670–5.

27. Bukalo O, Pinard CR, Silverstein S, Brehm C, Hartley ND, Whittle N, et al. Prefrontal inputs to the amygdala instruct fear extinction memory formation. Sci Adv. 2015;1(6).

28. Giustino TF, Maren S. The Role of the Medial Prefrontal Cortex in the Conditioning and Extinction of Fear. Front Behav Neurosci. 2015;9:298.

29. Kreutzmann JC, Jovanovic T, Fendt M. Infralimbic cortex activity is required for the expression but not the acquisition of conditioned safety. Psychopharmacology (Berl). 2020.

30. MacLeod JE, Bucci DJ. Contributions of the subregions of the medial prefrontal cortex to negative occasion setting. Behav Neurosci. 2010;124(3):321–8.

31. Meyer HC, Bucci DJ. The contribution of medial prefrontal cortical regions to conditioned inhibition. Behav Neurosci. 2014;128(6):644–53.

32. Apergis-Schoute AM, Gillan CM, Fineberg NA, Fernandez-Egea E, Sahakian BJ, Robbins TW. Neural basis of impaired safety signaling in Obsessive Compulsive Disorder. Proc Natl Acad Sci U S A. 2017;114(12):3216–21.

33. Greenberg T, Carlson JM, Cha J, Hajcak G, Mujica-Parodi LR. Ventromedial prefrontal cortex reactivity is altered in generalized anxiety disorder during fear generalization. Depress Anxiety. 2013;30(3):242–50.

34. Klavir O, Genud-Gabai R, Paz R. Functional connectivity between amygdala and cingulate cortex for adaptive aversive learning. Neuron. 2013;80(5):1290–300.

35. Stujenske JM, Likhtik E, Topiwala MA, Gordon JA. Fear and safety engage competing patterns of theta-gamma coupling in the basolateral amygdala. Neuron. 2014;83(4):919–33.

36. Cho JH, Deisseroth K, Bolshakov VY. Synaptic encoding of fear extinction in mPFC-amygdala circuits. Neuron. 2013;80(6):1491–507.

37. Pollak DD, Monje FJ, Lubec G. The learned safety paradigm as a mouse model for neuropsychiatric research. Nat Protoc. 2010;5(5):954–62.

38. Rodgers RJ, Boullier E, Chatzimichalaki P, Cooper GD, Shorten A. Contrasting phenotypes of C57BL/6JOlaHsd, 129S2/SvHsd and 129/SvEv mice in two exploration-based tests of anxiety-related behaviour. Physiol Behav. 2002;77(2-3):301–10.

39. Fendt M, Fanselow MS. The neuroanatomical and neurochemical basis of conditioned fear. Neurosci Biobehav Rev. 1999;23(5):743–60.

40. Fonio E, Benjamini Y, Golani I. Short and long term measures of anxiety exhibit opposite results. PLoS One. 2012;7(10):e48414.

41. McLean CP, Anderson ER. Brave men and timid women? A review of the gender differences in fear and anxiety. Clin Psychol Rev. 2009;29(6):496–505.

42. McLean CP, Asnaani A, Litz BT, Hofmann SG. Gender differences in anxiety disorders: prevalence, course of, illness, comorbidity and burden of illness. J Psychiatr Res. 2011;45(8):1027–35.

43. Nolen-Hoeksema S, Aldao A. Gender and age differences in emotion regulation strategies and their relationship to depressive symptoms. Personality and Individual Differences. 2011;51(6):704–8.

44. Dugas MJ, Freeston MH, Ladouceur R, Rheaume J, Provencher M, Boisvert JM. Worry themes in primary, GAD, secondary, GAD, and other anxiety disorders. J Anxiety Disord. 1998;12(3):253–61.

45. Aikins DE, Craske MG. Cognitive theories of generalized anxiety disorder. Psychiatr Clin North Am. 2001;24(1):57–74, vi.

46. Morriss J, Macdonald B, Van Reekum CM. What Is Going On Around Here? Intolerance of Uncertainty Predicts Threat Generalization. PLoS One. 2016;11(5):e0154494.

47. Chattarji S, Tomar A, Suvrathan A, Ghosh S, Rahman MM. Neighborhood matters: divergent patterns of stress-induced plasticity across the brain. Nat Neurosci. 2015;18(10):1364–75.

48. Weinberger NM, Miasnikov AA, Chen JC. The level of cholinergic nucleus basalis activation controls the specificity of auditory associative memory. Neurobiol Learn Mem. 2006;86(3):270–85.

49. Letzkus JJ, Wolff SB, Meyer EM, Tovote P, Courtin J, Herry C, et al. A disinhibitory microcircuit for associative fear learning in the auditory cortex. Nature. 2011;480(7377):331–5.

50. Aizenberg M, Mwilambwe-Tshilobo L, Briguglio JJ, Natan RG, Geffen MN. Bidirectional Regulation of Innate and Learned Behaviors That Rely on Frequency Discrimination by Cortical Inhibitory Neurons. PLoS Biol. 2015;13(12):e1002308.

51. Bishop SJ. Neurocognitive mechanisms of anxiety: an integrative account. Trends Cogn Sci. 2007;11(7):307–16.

52. Davis M, Walker DL, Miles L, Grillon C. Phasic vs sustained fear in rats and humans: role of the extended amygdala in fear vs anxiety. Neuropsychopharmacology. 2010;35(1):105–35.

53. Grillon C. Models and mechanisms of anxiety: evidence from startle studies. Psychopharmacology (Berl). 2008;199(3):421–37.

54. MacLeod C, Mathews A, Tata P. Attentional bias in emotional disorders. J Abnorm Psychol. 1986;95(1):15–20.

55. Cisler JM, Koster EH. Mechanisms of attentional biases towards threat in anxiety disorders: An integrative review. Clin Psychol Rev. 2010;30(2):203–16.

56. Sarter M, Bruno JP. Cognitive functions of cortical acetylcholine: toward a unifying hypothesis. Brain Res Brain Res Rev. 1997;23(1-2):28–46.

57. Sarter M, Bruno JP, Turchi J. Basal forebrain afferent projections modulating cortical, acetylcholine, attention, and implications for neuropsychiatric disorders. Ann N Y Acad Sci. 1999;877:368–82.

58. Hersman S, Cushman J, Lemelson N, Wassum K, Lotfipour S, Fanselow MS. Optogenetic excitation of cholinergic inputs to hippocampus primes future contextual fear associations. Sci Rep. 2017;7(1):2333.

59. Zaborszky L, Gombkoto P, Varsanyi P, Gielow MR, Poe G, Role LW, et al. Specific Basal Forebrain-Cortical Cholinergic Circuits Coordinate Cognitive Operations. J Neurosci. 2018;38(44):9446–58.

60. Guo W, Robert B, Polley DB. The Cholinergic Basal Forebrain Links Auditory Stimuli with Delayed Reinforcement to Support Learning. Neuron. 2019;103(6):1164–77 e6.

61. Concina G, Cambiaghi M, Renna A, Sacchetti B. Coherent Activity between the Prelimbic and Auditory Cortex in the Slow-Gamma Band Underlies Fear Discrimination. J Neurosci. 2018;38(39):8313–28.

62. Corches A, Hiroto A, Bailey TW, Speigel JH, 3rd, Pastore J, Mayford M, et al. Differential fear conditioning generates prefrontal neural ensembles of safety signals. Behav Brain Res. 2019;360:169–84.

63. Harrison BJ, Fullana MA, Via E, Soriano-Mas C, Vervliet B, Martinez-Zalacain I, et al. Human ventromedial prefrontal cortex and the positive affective processing of safety signals. Neuroimage. 2017;152:12–8.

64. Karalis N, Dejean C, Chaudun F, Khoder S, Rozeske RR, Wurtz H, et al. 4-Hz oscillations synchronize prefrontal-amygdala circuits during fear behavior. Nat Neurosci. 2016;19(4):605–12.

65. Taub AH, Perets R, Kahana E, Paz R. Oscillations Synchronize Amygdala-to-Prefrontal Primate Circuits during Aversive Learning. Neuron. 2018;97(2):291–8 e3.

66. Benjamini Y, Fonio E, Galili T, Havkin GZ, Golani I. Quantifying the buildup in extent and complexity of free exploration in mice. Proc Natl Acad Sci U S A. 2011;108 Suppl 3:15580–7.

